# Hybrid Clustering of single-cell gene expression and cell spatial information via integrated NMF and k-means

**DOI:** 10.1101/2020.11.15.383281

**Authors:** Sooyoun Oh, Haesun Park, Xiuwei Zhang

**Affiliations:** School of Computational Science and Engineering, Georgia Institute of Technology, Atlanta GA 30332, USA

## Abstract

**Motivation:** Recent advances in single cell transcriptomics have allowed us to examine the identity of single cells, which has led to the discovery of new cell types and high resolution maps of cell type composition in tissues. Technologies that measure multiple modalities of single cell data provide a more comprehensive picture of a cell, but they also create challenges for data integration tasks.

**Results:** In our work, we jointly consider the spatial location and gene expression profiles of cells to determine their identity. Specifically, we have developed scHybridNMF (single-cell Hybrid Nonnegative Matrix Factorization), which performs cell type identification by incorporating single cell gene expression data with cell location data. We combined nonnegative matrix factorization (NMF) with k-means clustering to cohesively represent high-dimensional gene expression data and low-dimensional location data, respectively. We show that scHybridNMF can utilize location data to improve cell type clustering. In particular, we show that under multiple scenarios, including the cases where there is a small number of genes profiled and the location data is noisy, scHybridNMF outperforms sparse NMF, k-means, and an existing method (HMRF) that also uses cell location and gene expression data for cell type identification.

**Availability:** https://github.com/soobleck/scHybridNMF

## 1 Introduction

Advances in single cell RNA-Sequencing (scRNA-Seq) technology have provided an unprecedented opportunity for researchers to study the identity and mechanisms of single cells (Morris, 2019). While scRNA-Seq data is a major type of data used to study single cells, it cannot fully determine the identity of a cell (McKinley *et al*., 2020). As such, it is important to consider other modalities such as chromatin accessibility (Cusanovich *et al*., 2015), protein abundance (Peterson *et al*., 2017), and spatial locations (Ståhl *et al*., 2016; Wang *et al*., 2018) of single cells.

With the availability of these data, we have entered the era of multi-modal single-cell -omics, and effective computational methods are crucial in integrating multi-modal data to learn a comprehensive picture of inter- and intra-cell processes (Efremova and Teichmann, 2020; Stuart and Satija, 2019). Spatial location data can provide important information on the cells’ micro-environment and allow researchers to study cell-cell interactions (Mayr *et al*., 2019). This is because cells at nearby locations tend to form the same cell type – daughter cells tend to keep the same cell type and similar location as their mother cell.

Considering both the gene expression and location data can lead to more accurate cell type identification. Technologies that measure the location and gene expression of the same set of cells often have to comprise on the number of genes measured (Zhu *et al*., 2018). Clustering cells using smaller gene expression profiles can be inaccurate, so incorporating the cell location data can improve its accuracy. However, reconciling single cell gene expression and location data for cell type identification is challenging because different data types can have differing scales, distributions, and types of noise (Efremova and Teichmann, 2020).

We introduce a matrix low-rank approximation scheme, scHybridNMF (single-cell Hybrid NMF), to perform cell clustering by jointly processing 2-dimensional cell location and gene expression data. Previously, Zhu *et al* developed a HMRF (Hidden Markov Random Field) model and showed that the spatial location of cells can contribute to cell type identification (Zhu *et al*., 2018). We, however, use a matrix low-rank approximation scheme because of the ease of preserving data characteristics through constraints and optimization terms. Crafting a loss-based minimization objective that bakes in these data characteristics maximally utilizes this information to jointly-cluster cells. We combined nonnegative matrix factorization with a k-means clustering scheme to cohesively represent high-dimensional gene expression data and low-dimensional location data, respectively.

Such joint-clustering methods based on matrix low-rank approximation have been used in other contexts, such as document clustering (Du *et al*., 2019) and. Additionally, promising NMF models have been developed for cell type identification for data ranging from just scRNA-Seq data to encompassing multiple modalities (Duren *et al*., 2018; Jin *et al*., 2020; Kotliar *et al*., 2019; Shao and Höfer, 2017; Welch *et al*., 2019). However, none of these methods incorporate cell locations. We compare our scHybridNMF model with both the standalone NMF and the k-means methods, as well as the HMRF method which uses spatial location information. We show that scHybridNMF is particularly advantageous in two application scenarios: to use when the number of genes with gene expression data is small, or and when the location data is noisy.

## 2 Methods

Matrix low-rank approximations assume that a matrix can be well-approximated as a product of lower-rank matrices. Many biological clustering frameworks are designed as matrix low-rank approximation schemes because they can easily incorporate prior biological knowledge and data constraints. Likewise, we formulate our multimodal clustering algorithm as a combination of multiple low-rank approximations. This formulation is designed to guide the gene expression-based clustering of cells with cell location clusters.

### 2.1 Review of Sparse NMF and K-Means Clustering

As part of our design, we incorporate sparse NMF and k-means clustering for incorporating gene expression and cell location data. We chose these methods because they can easily be formulated as matrix low-rank approximations, and creating an objective that incorporates both of these methods is intuitive. Additionally, the individual characteristics of each method strongly match the characteristic we wish to preserve in the data.

K-means clustering is an unsupervised learning algorithm that clusters data points by comparing pairwise distances, usually determined by the Euclidean distance metric. This metric naturally pairs with location-based data because it determines the similarity between points by how physically close they are. The matrix formulation for a Euclidean distance-based k-means objective for clustering *L* ∈ℝ^2×*n*^ is below.

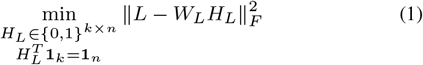

**1**_*k*_ and **1**_*n*_ are *k*-length and *n*-length vectors of all ones, respectively. The columns of *W*_*L*_ ∈ℝ^2×*k*^ contain *k* cluster centroids, and the columns of *H*_*L*_ ∈ℝ^*k*×*n*^ contain membership information for each data point. If a point *i* belongs to a cluster *j, H*_*L*_(*i, j*) = 1 and 0 for other clusters. The constraints preserve the hard-clustering quality of *H*_*L*_, as each data point can only belong to one cluster, which is equivalent to having one 1 per column of *H*_*L*_. Additionally, k-means clustering does not require any pre-processing on location data. Pre-processing input data may remove many of the underlying characteristics of the location data. As such, k-means clustering is a good fit for our two-dimensional location data.

NMF is a dimension reduction algorithm that is well-suited for high-dimensional data. It computes two nonnegative factors, *H*_*A*_ and *W*_*A*_ of a specified reduced dimension size *p*, of the nonnegative input matrix *A* ∈ℝ_+_^*m*×*n*^. *p* is generally much smaller than *m* and *n*. The columns of *W*_*A*_ ∈ℝ_+_^*m*×*p*^ contain *p* cluster representatives, and the columns of *H*_*A*_ ∈ℝ_+_^*p*×*n*^ contain the cluster membership information for each data point.

Sparse NMF, in particular, constrains the sparsity in each column of *H*_*A*_ to make it more suitable for hard-clustering (Kim and Park, 2007). It converts NMF, which can be interpreted as a soft-clustering method, into more of a hard-clustering method – a data point will have fewer nonzero entries in the cluster membership matrix, and therefore be represented by fewer cluster representatives. The results of sparse NMF may be interpreted as a hard clustering if we assign each data point to its respective maximal element in its column of *H*_*A*_. For example, if the largest element in the first column of *H*_*A*_ is in the second entry, we can interpret the first data point as belonging to the second cluster.

Below is the formulation for sparse NMF. The first term is the objective term for plain NMF factorization, which minimizes the difference between *A* and *W*_*A*_*H*_*A*_. Since *H*_*A*_ from NMF is not unique, we force the columns of *W*_*A*_ to have unit norm by normalizing the columns of the computed *W*_*A*_ and accordingly multiplying *H*_*A*_ by the reciprocal of the norm. The final term constrains the sparsity in each column of *H*_*A*_, and the second term limits the size of the elements in *W*_*A*_. This is to prevent the case that each element in *H*_*A*_ is minimized (and elements in *W*_*A*_ are maximized) while keeping a full sparsity pattern.

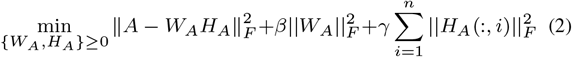

### 2.2 Multimodal Objective

Let *A* ∈ℝ_+_^*m*×*n*^ denote the gene expression matrix and *L* ∈ℝ^2×*n*^ denote the two-dimensional cell location coordinates, where *m* is the number of genes and *n* is the number of cells. To combine information about gene expression and spatial location, we compute *W*_*A*_ and *H*_*A*_ from sparse NMF and *W*_*L*_ and *H*_*L*_ from k-means clustering. To combine the clustering information from k-means clustering and NMF, we set *k* = *p*, which allows for a direct comparison between the two data. Additionally, we convert the *H*_*L*_ into a matrix of confidence scores, *Ĥ*_*L*_, to add consideration to how close each cell is to their cluster boundaries. We find the closest two cluster centroids, *b*_1_ and *b*_2_, to each cell *i*, then assign values to entries in *Ĥ*_*L*_ as in Eqn. (3). All other entries of *Ĥ*_*L*_ are zero.

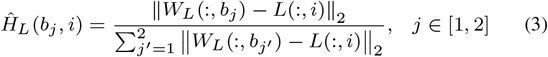

As such, we compare *H*_*A*_ with *Ĥ*_*L*_, and not with *H*_*L*_ directly. We use the following objective function for multimodal clustering:

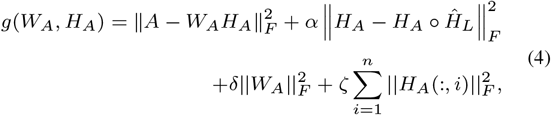

where º represents the element-wise product between two matrices. Every term but the second term in Eqn. (4) is from the sparse NMF objective in Eqn. (2). The second term combines NMF and k-means clustering results. Instead of forcing *H*_*A*_ and *Ĥ*_*L*_ to be similar overall, the second term forces *H*_*A*_ and *Ĥ*_*L*_ to be similar in terms of cluster memberships. In other words, we want the location of the largest element in each column of *H*_*A*_ and the location of the two nonzero elements in the corresponding column of *Ĥ*_*L*_ to match as much as possible.

The main focus of this work is to use cell location information to aid the gene expression-based clustering of cells. Because we are specifically adapting our gene clusters to incorporate location cluster information, our design seeks to align the cluster membership matrices found in both k-means and NMF while still considering the accuracy of the gene expression clustering. Because our method incorporates the predetermined location-based clusters, it would not make sense to add location clustering in the objective. That is why *Ĥ*_*L*_ but not the k-means objective, is in Eqn. (4).

### 2.3 Proposed Algorithm

We devise scHybridNMF to minimize Eqn. (4) using a consensus clustering on the clusters determined by sparse NMF on *A* and k-means on *L*. We set initial location cluster centroids for k-means clustering by computing the hard-cluster memberships of each cell in *H*_*A*_, then taking the mean of their locations. By design, there will be *k* different cluster centroids.

The crux of our algorithm is in the block coordinate descent for computing *H*_*A*_ and *W*_*A*_. These two terms are computed via an alternating nonnegative least squares (ANLS) formulation. We isolate the terms that involve *H*_*A*_ and *W*_*A*_ in Eqn. (4) to formulate the inputs into ANLS.

To solve for *H*_*A*_, we only need to combine the first and second terms in Eqn. (4). Given that the second term involves *H*_*A*_ twice, we reformulate the second term as follows:

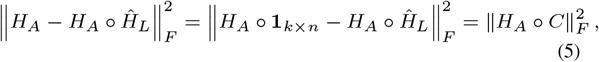

where *C* = 1_*k*×*n*_ − *Ĥ*_*L*_ and **1**_*k*×*n*_ the *k* × *n* matrix of all ones. We can represent an element-wise product in a block-ANLS formulation by computing the formulation column-by-column. Therefore, the new update rule for the first and second terms of Eqn. (4) is as follows:

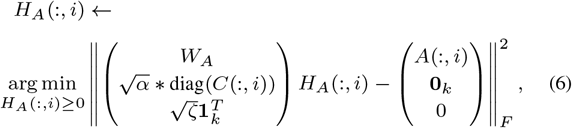

where *i* ∈ {1, …, *k*}, **1**_*k*_ is a *k*-length vector of all ones, and **0**_*k*_ is a *k*-length vector of all zeros. Each column in *H*_*A*_ is element-wise multiplied to each column in *C* in Eqn. (5), which can be represented as a left-multiplication of the column of *H*_*A*_ by a matrix whose diagonal entries are the corresponding column of *C*.

To solve for *W*_*A*_, we transposed the first and third terms in Eqn. (4):

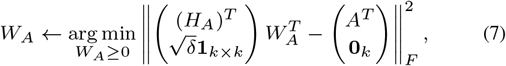

where **1**_*k*×*k*_ is a *k*-by-*k* matrix of all ones. The overall algorithm is described in Algorithm 1. There exist many stopping criteria that can ensure the proper convergence of our algorithm. We used the projected gradient, as used in SymNMF, to be the stopping criterion of scHybridNMF (Kuang *et al*., 2015).

#### Algorithm 1: scHybridNMF: an algorithm to minimize Eqn. (4)

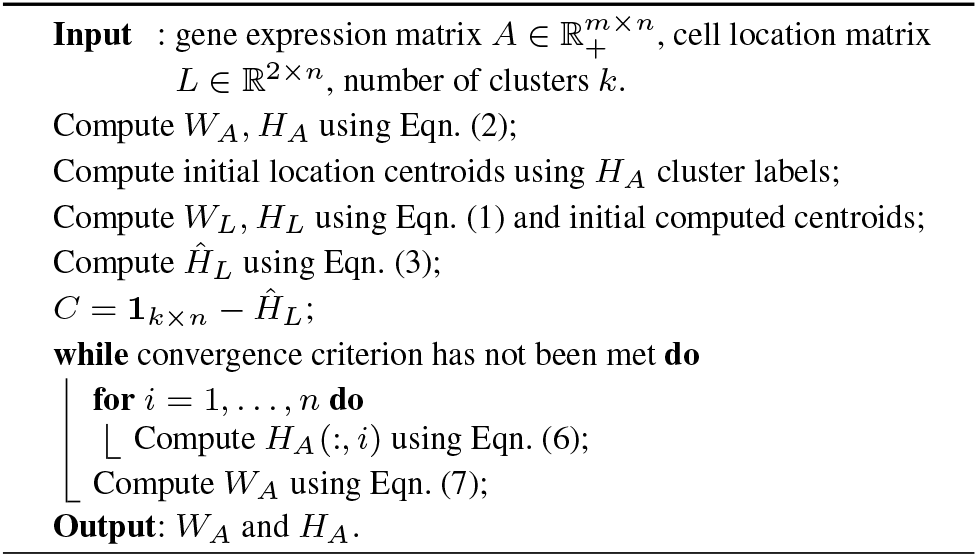

### 2.4 Convergence of Algorithm

We use a block coordinate descent (BCD) framework to optimize our objective function for clustering multimodal data. BCD solves subgroups of problems for each variable of interest, which iteratively minimizes the total objective function. Our objective aims to iteratively improve *W*_*A*_ and *H*_*A*_, which defines a two block coordinate descent framework. These comprise the minimization version of the two-block Gauss-Seidel method, which assigns *H*^(*j*)^and *W* ^(*j*)^ values that minimize a shared objective function, Eqn. (4), one-at-a-time.

An important theorem regarding general block Gauss-Seidel methods states that if a continuously differentiable function over a set of closed convex sets is minimized by block coordinate descent, every solution that uniquely minimizes the function in block coordinate descent is a stationary point (Bertsekas *et al*., 1997). This theorem has the additional property that the uniqueness of the minimum is not necessary for a two-block Gauss-Seidel nonlinear minimization scheme (Grippo and Sciandrone, 2000). This was used to show that a two-block formulation for solving plain NMF via alternating least squares guarantees convergence (Kim *et al*., 2014).

Given the constrained nonlinear minimization objective in Eqn. (4), we can rewrite the block coordinate descent as two ANLS formulations, which follow from Eqns. (6) and (7):

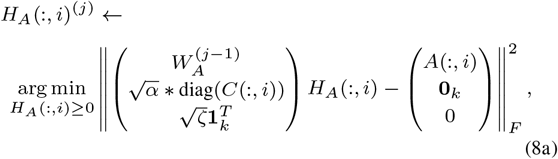

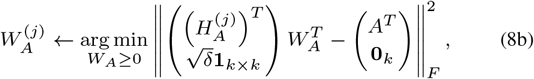

Eqns. (8a) and (8b) are executed consecutively to solve for *H*_*A*_ and *W*_*A*_. We consider Eqn. (8a) to be one block calculation because the calculations for each individual column are independent of each other. In other words, the calculation of a column of 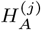 does not involve any other column of 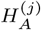. We are then able to apply this theorem because Eqns. (8a) and (8b) constitute a valid minimization scheme equivalent to minimizing Eqn. (4). As such, we get the following property, which guarantees the convergence of our algorithm:

Theorem 1. *Every limit point* 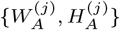 *calculated iteratively via Eqns. (8a) and (8b) is a stationary point of Eqn. (4)*.

## 3 Results

To prove the robustness of our algorithm, we tested it against simulated and real data. For real data, we tested the performance of our algorithm on the seqFISH dataset, which catalogues a mouse brain cortex. In both cases, we compare against HMRF, an existing method that also performs consensus cell clustering on gene expression and cell location data.

### 3.1 Simulated Data

We use SymSim (Zhang *et al*., 2019) to simulate single cell gene expression data where cells are from six cell types. Each dataset has 1600 cells and 600 genes. The number of genes is set to reflect the relatively low number of genes profiled in some spatially-resolved single cell gene expression datasets.

We simulate the location data for the cells in a 2-d space such that cells belonging in the same cell type are closely located in the 2-d location space. This procedure mimics the cell division process in a tissue. First, in the 2-d space, we choose a starting location for the earliest cell in each cell type. Then, for each cell type, a new cell is added in the following fashion: we randomly choose an existing cell of the same type to be its parent, and place the new cell next to the parent cell. If there is no available position next to the parent cell, then the new cell is put in a random empty position.

We consider different scenarios for the cell location data depending on how well the clusters are separated in the space. We denote clusters that are well separated as *w-separated*, and clusters that are not well separated as *n-separated*. For each of these scenarios, we generate location data with and without noise. In noisy data, cells from different cell types are mixed in the location space, and in data without noise, cells in the same cell type are all located together. We obtain the noisy location data from location data without noise by randomly choosing a percentage of cells and assigning them locations which are not in the main region of their original cell type. This is to more accurately emulate real-life data. Fig. 1 shows examples of these cases, where the case of n-separated with noise (as shown in Fig. 1d) is closest to real-life data.

**Fig. 1.**
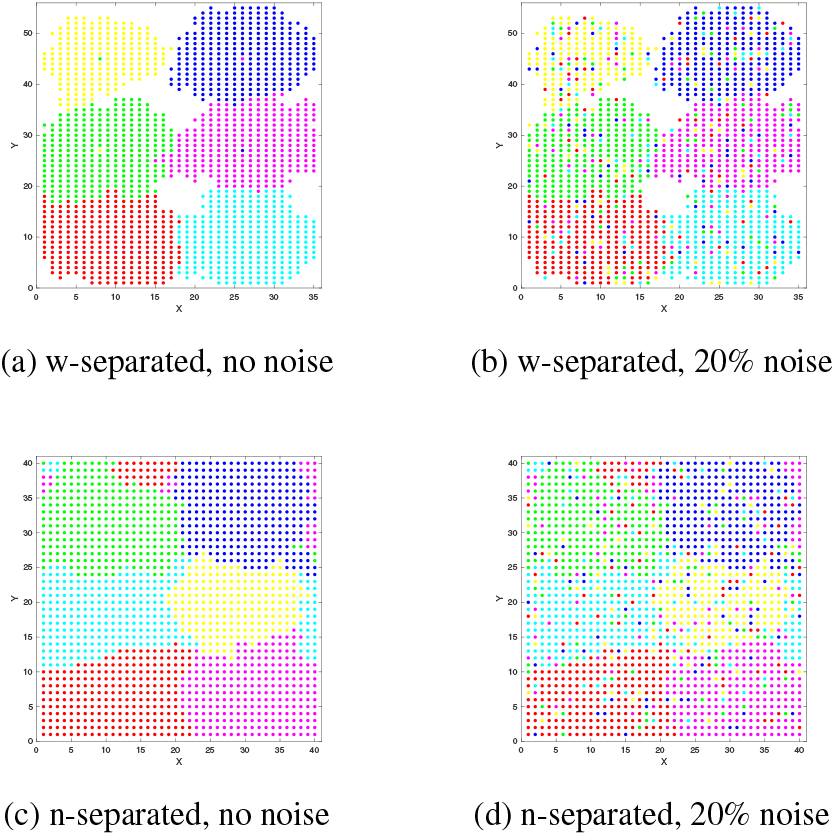
An example of noise in location data. The data had *σ* = 0.3 and 20% noise in location. In each plot, there are six point colors that correspond to the colorized true cluster labels. (a) and (b) share the same ground truth w-separated cluster labels, and (c) and (d) share the same ground truth n-separated cluster labels.

SymSim has a parameter *σ* (sigma) which adjusts the within-cluster heterogeneity. When *σ* increases, the gene expression-based clusters are less separable. In our experiments we test the performance of our algorithm with varying *σ*. The hypothesis is that when *σ* increases the data is more difficult for clustering algorithms using the gene expression alone, and we should gain more improvement through integrating location data.

In our tests, we use different values for *σ* to generate single cell gene expression data. For each parameter setting, 10 datesets are generated. To test on datasets where even less genes are measured, we randomly sample 50% of the genes from the original gene expression datasets to get a total of 300 genes. In total, we tested against noisy location data, which are w-separated and n-separated, paired with gene expression data with and without sampling. For w-separated location data, we use *σ* = {0.4, 0.5, 0.6, 0.7}, and for n-separated location data, we use *σ* = {0.3, 0.4, 0.5, 0.6}. We are using lower *σ* for n-separated data because we wanted to analyze the performance of our algorithm on cases where gene expression data may be much more useful than location data in determining cell identity. This is a boundary case that showcases the ability of our algorithm to balance the influence of cell locations to the consensus clustering.

All of the parameters in Eqns. (2) and (4) have an impact on the results, and we provide analytical forms for setting them. We keep *β* and *δ* equal and constant, at 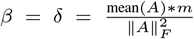. We also keep 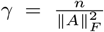. To balance the influence of the *H*_*A*_ sparsity term with respect to the rest of Eqn. (4), we set 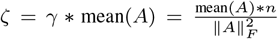. Finally, we determined 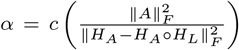, where *c* is a user-input term whose recommended range is between 0 and 10. In our experiments, we use the same *c*-values for data with and without sampling. For w-separated data, we use *c* = [3.5, 4, 4.5, 7.5], and for n-separated data, we use *c* = [1.5, 4, 3.5, 4]. Note that *H*_*L*_ is calculated from k-means clustering directly (and is not *Ĥ*_*L*_). We used this formulation for *α* because it normalises the second term by its worst possible value for 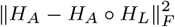 (as per Thm. 1, every iteration improves this norm difference). It also accounts for the normalising term 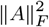 for the first term.

We compared the quality of clusters determined by gene expression clustering, cell location clustering, and hybrid clustering of both data. The methods we used for gene expression clustering were sparse NMF and PCA plus k-means clustering, which provides a baseline for the performance of sparse NMF – in cases of higher sigma values, a poor performance from PCA plus k-means clustering justifies the lower performance of sparse NMF. For location-based clustering, we just used k-means clustering. To cluster the combination of the two data, we used scHybridNMF and HMRF, one of the only published algorithms that also performs a consensus clustering of the two data.

As an example, Fig. 2 shows the tSNE plots, which visualize high-dimensional data, of the gene expression data clusters produced by NMF, scHybridNMF, and the ground truth labels. This shows that our method improves the performance of cell clustering of the gene expression immensely.

**Fig. 2.**
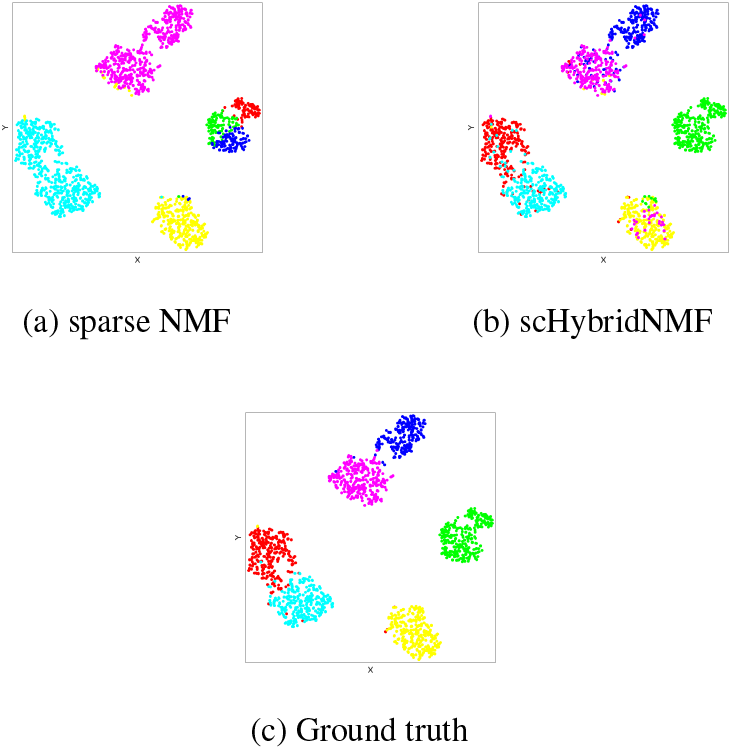
An example of NMF vs scHybridNMF vs ground truth clusters for gene expression. The data had *σ* = 0.3, 30% of the genes sampled, 20% noise in location, and w-separated location data. In each plot, there are six point colors that correspond to the six cluster labels.

To quantitatively evaluate the performance of scHybridNMF on our data, we calculated the adjusted Rand index (ARI) between the calculated clusters and the ground truth clusters for each set of experiments. In this context, ARI quantifies how similar two clusterings are to each other while correcting for chance. If the ARI of a clustering is very similar to the ground truth clustering, the ARI value should be close to 1. To ensure that there was an even comparison between sparse NMF, k-means clustering, and scHybridNMF, we calculated the sparse NMF and k-means clustering ARIs for the clusters that were used as steps 1 and 2 in Algorithm 1.

#### 3.1.1 Experiment 1: All Genes, Noisy Location Data

We consider the case where there is noise in the location data, but no gene sampling. A dataset with many genes tests how well scHybridNMF can discern between important differences in gene expression to determine cell type. We used different amounts of noise for different data, with w-separated data having 20% and 30% noise and n-separated data having 10% and 20% noise. This is to account for the inherent ease in location-based clustering for well-separated data in contrast to not well-separated data. For each location data with no noise, we generated 10 noisy location datasets and calculated the average ARI over 100 location-gene expression pairs, which accounts for each *σ* and noise percentage. For HMRF, we sampled 4 matrices from each *σ* value (2 for each location noise percentage) and averaged the performance of HMRF across 3 values (25, 50, 75) for a parameter that account for smoothness. We plotted the average values as a function of *σ* in Figs. 3 and 4.

**Fig. 3.**
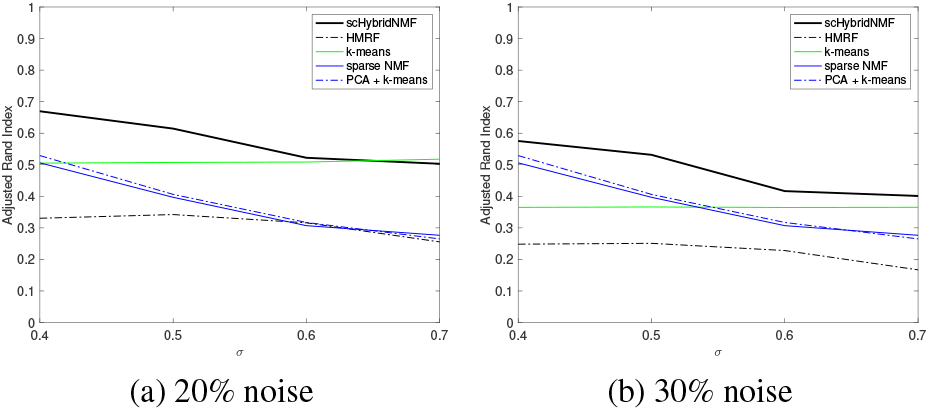
Using all genes with noisy data on w-separated data.

**Fig. 4.**
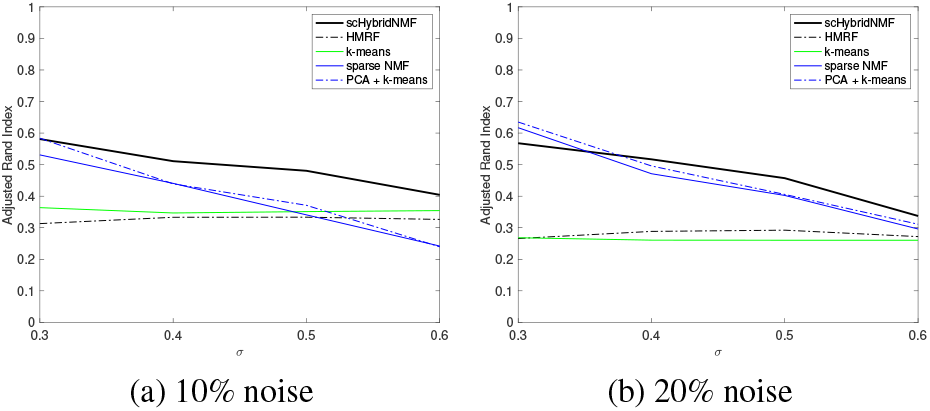
Using all genes with noisy data on n-separated data.

In Figs. 3 and 4, our algorithm had higher ARI values than every other case. This is especially relevant in Figs. 3, where the performance of k-means clustering often outperforms all of the purely gene expression-based methods. Improving over cases where k-means performs well shows that our algorithm can balance gene expression and spatial location when location data is heavily favored. Even with the decreasing performance of the k-means clustering results, scHybridNMF improves tremendously over sparse NMF. This is especially evident in Fig. 3, where scHybridNMF achieves a much higher performance than both sparse NMF and k-means, indicating that scHybridNMF is able to gather useful information from both standalone methods, and that it has high potential to be successful on real-world data.

#### 3.1.2 Experiment 2: Sampled Genes, Noisy Location Data

In our other case, we investigated the scenario of noisy location data and a small number of genes, which is the most challenging scenario. This resembles real-world data the closest because of the limitations of current sequencing technology. This is because many current technologies that pairwise measure the gene expression and spatial locations of single cells cannot also sequence many genes (Zhu *et al*., 2018).

To get a small number of genes, we randomly sampled 50% of the genes to get 300 randomly sampled genes. Over each *σ*-value, we calculated 5 random gene samples over 10 location noise randomizations (for each noise percentage) for each of the 10 location-gene expression pairs. Again, we sample 4 location-gene expression data pairs from each sigma value (with 2 for each noise percentage) and run HMRF on them. We take the average performance across 3 smoothness parameter values. We plotted the average ARI values for sampling 50% of the genes as a function of *σ* in Figs. 5 and 6. In this experiment, we also run the existing HMRF (Zhu *et al*., 2018) method which also performs cell clustering using both cell location and gene expression data, on the same datasets.

**Fig. 5.**
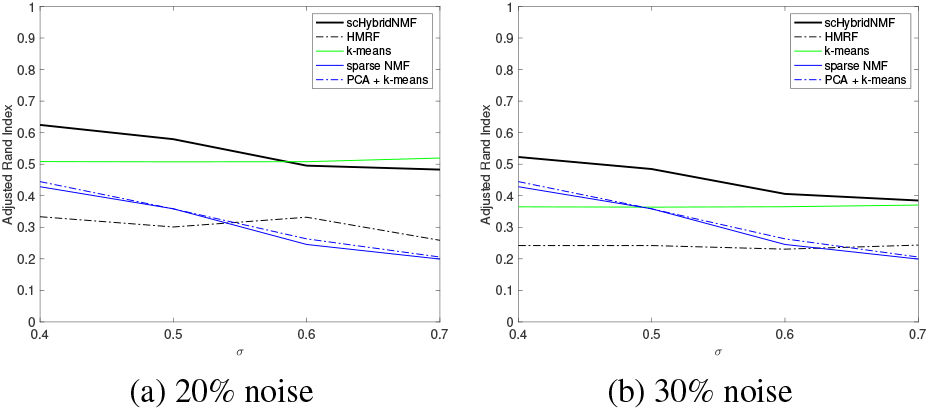
Using 50% of the genes with 20% and 30% noise in w-separated data.

**Fig. 6.**
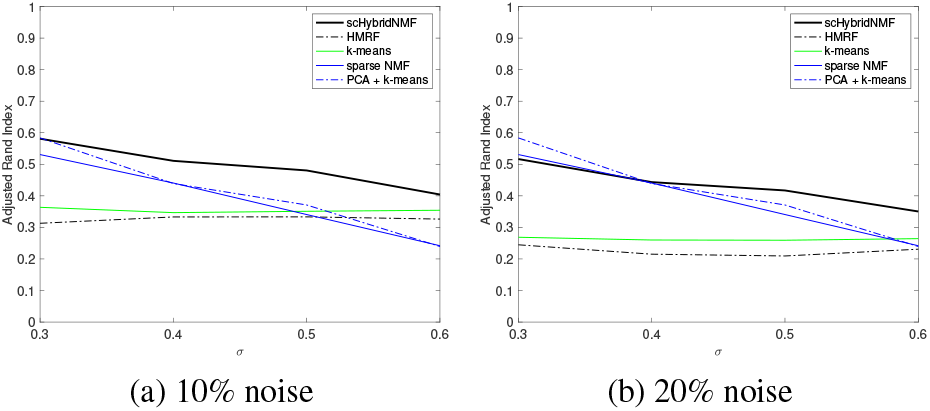
Using 50% of the genes with 10% and 20% noise in n-separated data.

The results show a clear distinction between scHybridNMF and NMF and k-means clustering. For the w-separated location data, the ARI values of scHybridNMF significantly exceeds those of NMF and HMRF clustering. In data with n-separated location data, scHybridNMF tends to outperform both k-means clustering and NMF, and ourperforms HMRF in all cases. Considering the fact that we used *c*-values that did not vary very much across the experiments, it is impressive how the performance of scHybridNMF remained consistently high.

These experiments show that scHybridNMF is robust to small datasets with noisy locations and a subset of the total number of genes. This sort of data is prevalent in the real world, and the fact that our algorithm performs the strongest relative to individually using NMF or k-means on this data indicates that it is likely to be successful for real data.

### 3.2 Real Data

We uses mouse brain cortex gene expression and location data from another consensus clustering scheme, Giotto, which utilizes the HMRF algorithm (Dries *et al*., 2019). This data has been adapted from the seqFISH+ dataset (Eng *et al*., 2019), and it has been annotated with the locations of specific cells as well as their gene expression levels. To examine the regions that have varied genetic expressions, we isolated the 523 cortex cells and filtered the genes to keep those with mean greater than 0.7 and correlation of variation greater than 1.2 measured across all cells. The analysis from (Dries *et al*., 2019) indicates that there may be 9 clusters, so to be able to compare with their results, we also set the number of clusters *k* = 9. We also set *c* = 4 10^*-*13^ and 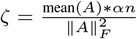. All other parameters kept their analytic formulations used in simulated data.

As in Fig. 7, we get 9 distinct clusters, which can correspond to the spatial domains in mouse brain, like those found in (Dries *et al*., 2019). In the cortex of a brain, we have 6 layers. In numerical order, they are the molecular, external granular, pyramidal, inner granular, ganglionic, and multiform layers. Given the k-means clusters in Fig. 7, it is obvious that purely location-based clustering cannot capture the true shape of the anatomical layers. As such, we need to combine both the gene expression and location data to accurately model real-world data.

**Fig. 7.**
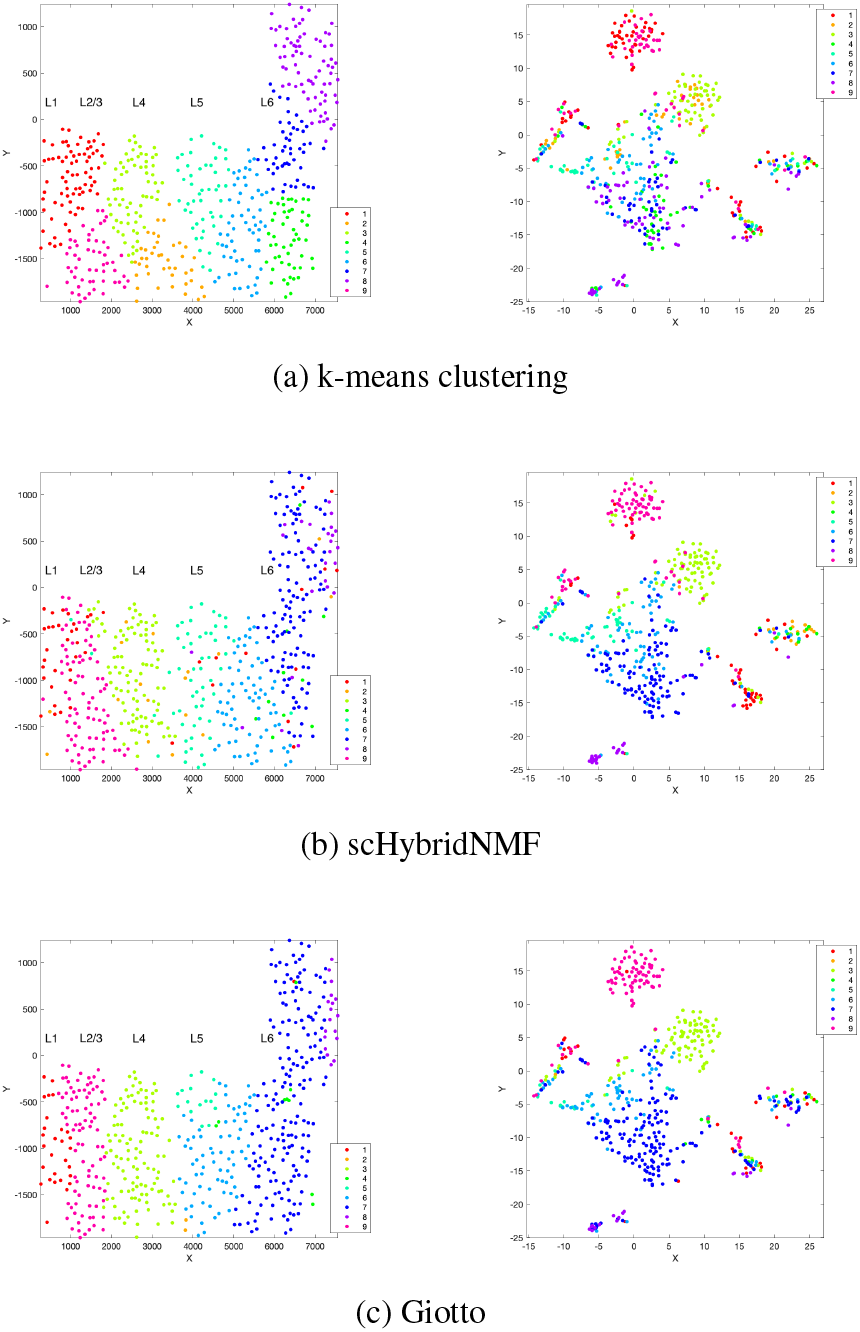
The location-based k-means clustering results and consensus clustering results of scHybridNMF and Giotto on seqFISH data. In each plot, there are nine point colors that correspond to the nine cluster labels. The left-hand plots represent the cells’ locations, and the right-hand plots show tSNE-reduced gene expression matrices. Layers 1-6 of the cortex are labeled with L1-L6, as in Dries et al. (2019).

The clusters determined by scHybridNMF are consistent with the underlying anatomical layers of the brain cortex. For example, cluster 1 directly matched up with layer 1. For the most part, the clusters determined by Giotto resemble the layers as well. However, our cluster 5 in Fig. 7 has a more rectangular shape than the small circular cluster 5 in Giotto.

This strongly suggests that our clusters, which are contiguous rectangular regions, resemble the anatomical layers of the mouse brain cortex more accurately than the Giotto clusters, which are more amorphous.

We further investigated clusters 5 and 6 assigned by scHybridNMF and Giotto to see which method provided more biologically meaningful results. We computed differentially-expressed genes between the two clusters by performing a *t*-test for each of the *m* candidate genes’ expression levels in the cells of the two clusters. We chose the top 50 genes with the smallest p-values, and performed a gene ontology (GO) analysis to find which functions the selected genes were enriched in. The GO analysis was performed with DAVID, an online functional annotation tool (Huang *et al*., 2009a,b). We aimed to record the top 10 GO terms with the most significant p-values, all of which were required to be below 0.05. However, there were only 5 GO terms for the Giotto cluster-derived genes. The top 10 GO terms obtained with scHybridNMF cluster labels are in Table 1, while Table 2 shows the top 5 GO terms obtained with the Giotto clustering.

**Table 1.** Top 10 GO terms determined from top 50 differentially-expressed genes between clusters 5 and 6 in scHybridNMF.

| Category | Term | P-Value |
| --- | --- | --- |
| GOTERM_CC | Dendrite | 7.3E-4 |
| GOTERM_BP | Negative Regulation of Cell Cycle | 3.6E-3 |
| GOTERM_BP | Neuron Differentiation | 3.9E-3 |
| GOTERM_CC | Myelin Sheath | 9.0E-3 |
| GOTERM_MF | Ion Channel Inhibitor Activity | 1.3E-2 |
| GOTERM_CC | Neurofilament | 2.0E-2 |
| GOTERM_BP | Neurofilament Cytoskeleton Organization | 2.5E-2 |
| GOTERM_CC | Synapse | 2.6E-2 |
| GOTERM_BP | Exocytosis | 2.8E-2 |
| GOTERM_BP | Locomotory Behavior | 3.0E-2 |

**Table 2.** Top 5 GO terms determined from top differentially-expressed genes between clusters 5 and 6 in Giotto.

| Category | Term | P-Value |
| --- | --- | --- |
| GOTERM_CC | Dendrite | 9.3E-4 |
| GOTERM_BP | Regulation of Adenylate Cyclase Activity | 9.7E-3 |
| GOTERM_BP | Regulation of cAMP-Mediated Signaling | 1.2E-2 |
| GOTERM_BP | Cytoskeleton Organization | 2.6E-2 |
| GOTERM_CC | Neuronal Cell Body | 3.6E-2 |

We observe that, in Table 1, we have 6 terms specific to neuronal cells, while Table 2 has only 2 terms related to neuronal cells. The fact that many highly differential genes from scHybridNMF are related to fundamental properties of neuronal cells specifically, we can assume that the two clusters identified by scHybridNMF have unique cellular identities. Since the clusters occupy two cortex layers, which are comprised of different neuron types, it would make sense to separate them based on neuronal differences. As we cannot say the same about the GO profile of the differential genes of Giotto, this gives additional credence to the accuracy of our cluster boundary between clusters 5 and 6. Furthermore, most of Giotto’s top GO terms have larger p-values than scHybridNMF’s top GO terms. This indicates that the clusters determined by scHybridNMF are more likely to be biologically meaningful than those discovered by Giotto.

## 4 Conclusions and Discussions

In this paper, we presented a hybrid clustering approach that can better identify cell types by incorporating the strengths of sparse NMF and k-means clustering, which work well on high-dimensional single cell gene expression data and low-dimensional location data. We demonstrated the robustness of our algorithm through testing it on different simulated data configurations as well as on a real mouse brain cortex dataset.

We show that our hybrid framework, scHybridNMF, significantly improves over the clustering accuracy of using sparse NMF alone on gene expression data by integrating location information. This is particularly useful for the cases where sparse NMF performance is affected by a low number of genes in the gene expression data or high within-cluster heterogeneity. scHybridNMF also outperforms k-means clustering with only location data under realistic scenarios. Through combining two classical methods for clustering, sparse NMF and k-means, scHybridNMF can exploit both the high and low dimensional data and achieve better performance than using either of the standalone methods, as well as an existing method HMRF. We also observed that our algorithm found biologically-meaningful clusters within real data. Against the performance of Giotto, which uses HMRF in its consensus clustering, our algorithm more successfully recreated a biologically meaningful separation between cells near layers 5 and 6 of the cortex.

This framework is inherently flexible, owing to its simple matrix low-rank approximation formulation. As such, it can be extended via additional matrix terms and constraints to include more types of data or to perform biclustering. For example, we can include potential gene-gene interaction data to perform co-clustering of both cells and genes. The inferred gene clusters can be further used to study regulatory mechanisms in the cells and reconstruct gene regulatory networks.

## Acknowledgements

We thank our colleagues for their editorial comments and discussions. This work was supported in part by the US National Science Foundation DBI-2019771. Any opinions, findings and conclusions or recommendations expressed in this material are those of the authors and do not necessarily reflect the views of NSF.

